# Conformational dynamics of an intrinsically disordered receptor enable signal transduction

**DOI:** 10.64898/2026.09.02.748842

**Authors:** Yuma Myokan, Huazi Zhang, Tsunaki Hongu, Keisuke Sato, Sarenqiqige, Katsuya Sakai, Yilmaz Neval, Kunio Matsumoto, Takashi Sumikama, Mikihiro Shibata, Noriko Gotoh

## Abstract

Signal transduction is initiated when activated cell-surface receptors physically engage intracellular signaling proteins. Although structural and biochemical studies have established the molecular architecture of receptor tyrosine kinase (RTK) signaling, the earliest dynamic events that couple receptor activation to intracellular signal propagation have remained inaccessible because existing approaches provide static structural snapshots or ensemble-averaged measurements rather than direct observation of molecular motion. Here, using high-speed atomic force microscopy (HS-AFM), we directly visualize these initial molecular events at the single-molecule level in real time. Using fibroblast growth factor receptor 1 (FGFR1) as a model, we show that receptor autophosphorylation releases its intrinsically disordered juxtamembrane region, enabling extensive conformational dynamics of the kinase domain that permit productive engagement of the signaling adaptor FRS2α, a key initiator of downstream RTK signaling. In contrast, a kinase-inactive mutation or the clinically approved FGFR inhibitor futibatinib restricts receptor motion, stabilizes a compact receptor configuration, and markedly reduces adaptor engagement. Molecular dynamics simulations suggest that the disruption of inhibitory interactions between the juxtamembrane region and the kinase domain provides a structural explanation for the observed receptor dynamics. Together, our findings uncover a previously inaccessible physical mechanism underlying the initiation of RTK signaling and establish HS-AFM as a powerful platform for visualizing receptor dynamics, providing a framework for mechanistic studies of signal transduction and for the development and mechanistic evaluation of compounds that modulate receptor signaling.

## Introduction

Cells continuously sense and respond to their environment by transmitting information across the plasma membrane. Receptor tyrosine kinases (RTKs) are central regulators of this process, coupling extracellular stimuli to intracellular signaling networks that govern development, tissue homeostasis and cancer ^1,2^. Over the past several decades, structural, biochemical and biophysical studies have established the molecular architecture of RTKs and the mechanisms of receptor activation, providing the foundation for the development of numerous clinically successful tyrosine kinase inhibitors ^3, 4,5,6^. Yet one fundamental question has remained unresolved: how is receptor activation physically converted into intracellular signal transduction?

Addressing this question has proven challenging because the earliest intracellular signaling events are mediated by intrinsically disordered receptor regions that undergo continuous conformational fluctuations rather than adopting stable three-dimensional structures ^7,8,9,10,11^. X-ray crystallography, cryo-electron microscopy and related structural approaches have transformed our understanding of receptor architecture and activation mechanisms, whereas biochemical studies have defined the molecular interactions and signaling pathways that follow receptor activation ^1,12,13,14^. Complementary biophysical approaches, including nuclear magnetic resonance spectroscopy, have further provided important insights into the conformational properties of intrinsically disordered proteins ^8^. However, direct visualization of the real-time molecular behavior through which these dynamic receptor regions physically engage intracellular signaling proteins has remained difficult.

Fibroblast growth factor receptor 1 (FGFR1) provides an excellent model with which to address this question ^15,16^. FGFR substrate 2α (FRS2α) is the principal signaling adaptor of FGFR1^12,14,17,18,19^. Its phosphotyrosine-binding (PTB) domain recognizes the juxtamembrane (JM) region of FGFR1, and receptor activation promotes FRS2α phosphorylation, thereby initiating multiple downstream signaling pathways, including the RAS–MAPK and PI3K–AKT cascades. Although the molecular components of this signaling axis, including PTB-domain recognition and FRS2α phosphorylation, have been extensively characterized, the physical mechanism by which receptor activation enables productive adaptor engagement has remains unresolved. High-speed atomic force microscopy (HS-AFM) is uniquely suited to addressing this question: it directly visualizes individual biomolecules in aqueous solution at nanometer spatial and sub-second temporal resolution, capturing molecular motion in real time at the single-molecule level ^20,21,22^.

Here, by combining HS-AFM, molecular dynamics simulations, biochemical analyses and pharmacological perturbation, we directly visualize the earliest molecular events that physically couple receptor activation to adaptor engagement. We show that autophosphorylation releases the intrinsically disordered juxtamembrane (JM) region of FGFR1 from the kinase domain, allowing the receptor to undergo large-scale conformational fluctuations that permit productive engagement of FRS2α. Conversely, a kinase-inactive mutation abolishes both receptor motion and adaptor binding, and the clinically approved FGFR inhibitor futibatinib ^23^ similarly restrict receptor motion. These findings reveal that kinase-dependent dynamics of an intrinsically disordered receptor region govern the initiation of RTK signaling.

## Results

### Single-molecule imaging reveals dynamic intracellular behavior of activated FGFR1

To determine whether the earliest intracellular events of RTK signaling can be visualized directly at the single-molecule level, we established a membrane-reconstituted imaging system for human FGFR1. Upon activation, FGFR1 undergoes kinase activation and autophosphorylation, leading to phosphorylation of its principal signaling adaptor FRS2α and activation of downstream signaling pathways, including the RAS–MAPK and PI3K– AKT pathways ^12,17,19^ (Fig. 1a). To directly investigate intracellular receptor dynamics independently of extracellular ligand-binding events, we generated a construct comprising the transmembrane domain (TM) together with the complete intracellular (IC) region of FGFR1 (FGFR1 TM/IC; residues 377–822), including the JM, kinase and C-terminal regions (Fig. 1b,c).

**Fig. 1.**
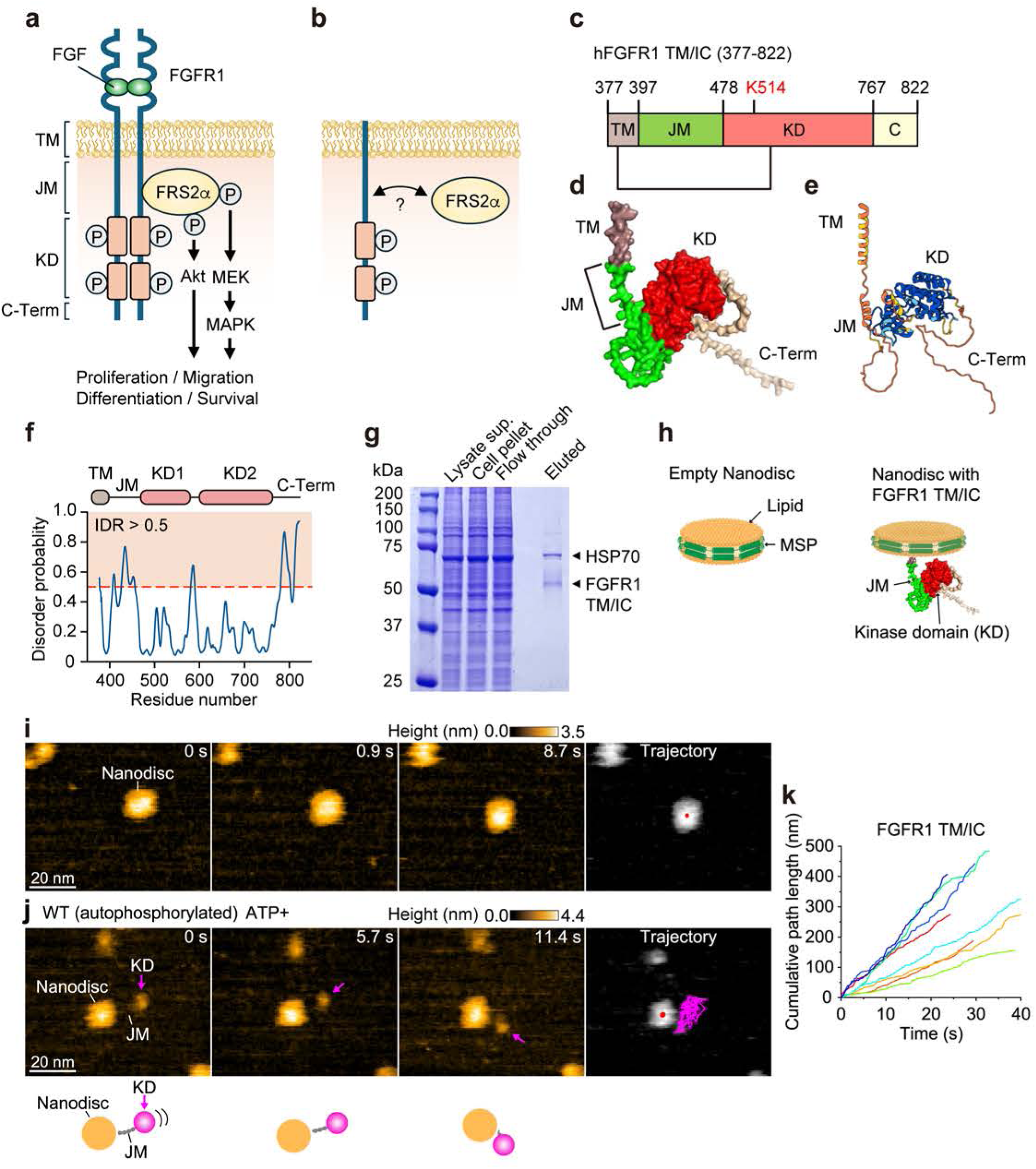
Single-molecule imaging reveals the conformational dynamics of activated FGFR1. **a,** Schematic overview of FGFR1 signaling. **b,** The earliest molecular events by which activated FGFR1 physically engages FRS2α remain unresolved. **c,** Domain organization of human FGFR1 TM/IC (residues 377–822), consisting of the transmembrane (TM), juxtamembrane (JM), kinase domain (KD), and C-terminal (C) region. The catalytically important lysine residue (K514) is indicated. **d,e,** AlphaFold3-predicted structure of FGFR1 TM/IC shown as surface (**d**) and ribbon (**e**) representations. **f,** Intrinsic disorder prediction identifies the JM and C-terminal regions as intrinsically disordered region (IDR) by local distance difference test (pLDDT). The dashed red line indicates the disorder threshold (IDR score = 0.5). **g,** Purified FGFR1 TM/IC and associated HSP70 are visualized by Coomassie blue staining. **h,** Schematic of empty nanodiscs and membrane-reconstituted FGFR1 TM/IC. **i,j,** Representative HS-AFM image sequences of empty nanodiscs (**i**) and membrane-reconstituted wild-type FGFR1 (**j**). Arrows indicate the kinase domain (KD). Schematic models corresponding to each image are shown below. The rightmost panels show trajectories of the center of the nanodisc (red point in gray nanodisc) and of the KD (magenta) tracked over approximately 30 s. Scale bars, 20 nm. Height scales are shown above each image series. **k,** Quantification of KD trajectories during HS-AFM imaging.

AlphaFold3 structural prediction ^24^ together with intrinsic disorder analysis predicted that the JM and C-terminal regions lack stable secondary structure, consistent with their classification as intrinsically disordered regions (Fig. 1d–f). FGFR1 TM/IC was expressed in mammalian cells and purified by FLAG affinity chromatography. Mass spectrometry identified FGFR1 as the predominant purified protein together with heat shock protein (HSP)70, a molecular chaperone associated with FGFR1 folding (Extended Data Fig. 1), and Coomassie blue staining confirmed proteins of the expected molecular masses (Fig. 1g).

To recreate a membrane environment suitable for single-molecule imaging, purified FGFR1 TM/IC was incorporated into lipid nanodiscs through its transmembrane domain (Fig. 1h). HS-AFM imaging of empty nanodiscs revealed particles smaller than 20 nm in diameter (Fig. 1i and Supplementary Movie 1). In contrast, membrane-reconstituted FGFR1 displayed an additional globular protrusion connected to the nanodisc by a thin, flexible linker, corresponding to the intracellular region (Fig. 1i). Following ATP-dependent receptor autophosphorylation, real-time HS-AFM imaging revealed extensive motion of the kinase domain around the nanodisc (Fig. 1j,k and Supplementary Movie 2).

These observations demonstrate that HS-AFM directly visualizes the dynamic intracellular behavior of activated FGFR1 at the single-molecule level and establish a platform for investigating how receptor conformational dynamics contribute to intracellular signal transduction. This experimental system enabled us to ask whether these kinase-dependent receptor dynamics permit productive engagement of the principal signaling adaptor FRS2α.

### FGFR1 conformational dynamics permit productive engagement of its signaling adaptor FRS2α

Having established an experimental platform for visualizing the intracellular dynamics of activated FGFR1, we next asked whether these receptor conformational dynamics permit productive engagement of its principal signaling adaptor, FRS2α. FRS2α contains an N-terminal myristoylation sequence, a phosphotyrosine-binding (PTB) domain that recognizes the FGFR1 juxtamembrane (JM) region, and a C-terminal region containing multiple tyrosine residues that recruit downstream signaling proteins following phosphorylation^17^ (Fig. 2a). AlphaFold3 prediction together with intrinsic disorder analysis indicated that the PTB domain adopts a compact folded structure, whereas most of the C-terminal region is intrinsically disordered (Fig. 2b,c).

**Fig. 2.**
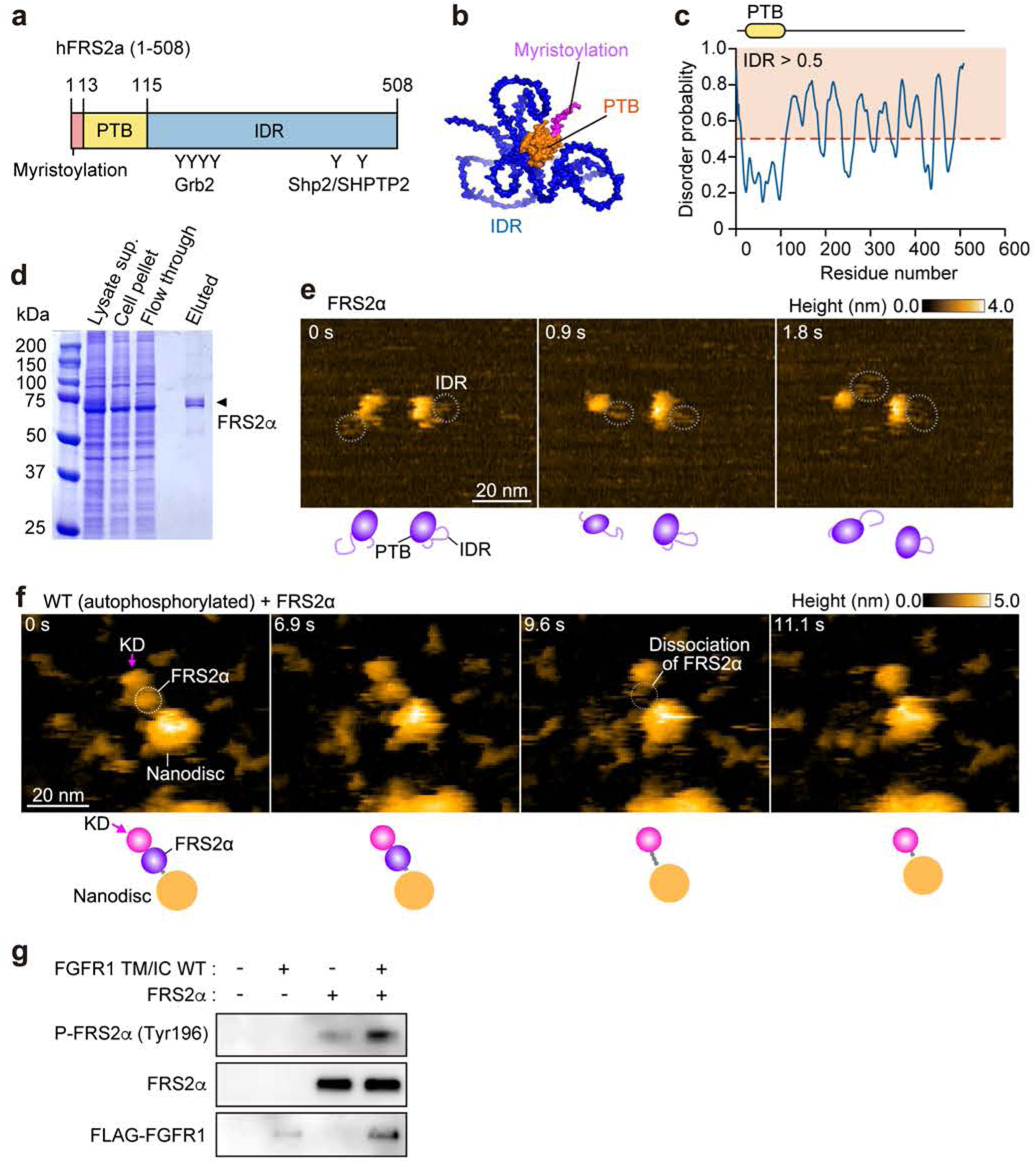
FGFR1 conformational dynamics enable productive engagement of the principal signaling adaptor FRS2α. **a,** Domain organization of human FRS2α, containing N-terminal myristoylation site, phosphotyrosine-binding (PTB) domain and IDR which carries tyrosine phosphorylation sites for Grb2 and Shp2/SHPTP2 recruitment. **b,** AlphaFold3-predicted structure of FRS2α. **c,** Intrinsic disorder prediction of FRS2α. The dashed red line represents the disorder threshold (IDR score = 0.5). **d,** Purified FRS2α is visualized by Coomassie blue staining. **e,** Representative HS-AFM image sequence of FRS2α. Dashed circles indicate the flexible IDRs. Schematic models corresponding to each image are shown below. **f,** Representative HS-AFM image sequence showing membrane-reconstituted wild-type (WT) FGFR1 engaging FRS2α. Arrows indicate the kinase domain. Dashed circles indicate FRS2α positioned adjacent to the FGFR1 juxtamembrane region. FRS2α dissociated from the complex at approximately 9.6 s (the second image from the right). Corresponding schematic models are shown below. **e,f**, Scale bars, 20 nm. Height scales are shown above each image series. **g,** In vitro phosphorylation of FRS2α (7.5 ng/μl) by purified WT FGFR1 (20 ng/μl).

To facilitate purification, Glycine (G) 2 was replaced by alanine to prevent N-terminal myristoylation without affecting the PTB domain (FRS2α G2A)^19^. Purified FRS2α(G2A) migrated at the expected molecular mass by Coomassie blue staining (Fig. 2d). HS-AFM imaging revealed a compact globular density corresponding to the PTB domain together with a highly flexible C-terminal region exhibiting extensive motion, consistent with its predicted intrinsically disordered nature (Fig. 2e and Supplementary Movie 3).

We next examined whether membrane-reconstituted FGFR1 could engage FRS2α under kinase-active conditions. HS-AFM revealed an additional globular density positioned between the nanodisc and the FGFR1 kinase domain, consistent with localization of the FRS2α PTB domain at its established binding site within the FGFR1 JM region (Fig. 2f and Supplementary Movie 4). The observed configuration demonstrates that the dynamic intracellular architecture of activated FGFR1 is compatible with productive adaptor engagement.

To determine whether adaptor engagement is accompanied by downstream signaling, purified FGFR1 TM/IC and FRS2α were subjected to an in vitro kinase assay. FGFR1 efficiently phosphorylated FRS2α at Tyr196 (Fig. 2g), confirming that the observed receptor–adaptor configuration is signaling competent.

Together, these findings indicate that kinase-active FGFR1 adopts a dynamic intracellular configuration that permits productive engagement and phosphorylation of its principal signaling adaptor. These observations prompted us to investigate whether this dynamic receptor state is governed by kinase activity.

### Kinase activity governs receptor conformational dynamics and productive adaptor engagement

Having established that activated FGFR1 adopts a dynamic intracellular configuration compatible with productive adaptor engagement, we next investigated whether these conformational dynamics are regulated by receptor kinase activity. We first examined a kinase-inactive mutant (K514A), in which the catalytic lysine (K) within the kinase domain is replaced by alanine (A)^25^ (Fig. 3a). Co-immunoprecipitation of FLAG-tagged FRS2α with wild-type or K514A FGFR1 TM/IC expressed in HEK293 cells revealed markedly reduced association of FRS2α with the kinase-inactive mutant (Fig. 3a). Purified K514A FGFR1 TM/IC was obtained using the same expression and purification strategy as wild-type FGFR1 (Fig. 3b), and loss of activation-loop phosphorylation at Tyr653 and Tyr654 confirmed disruption of kinase activity (Fig. 3c).

**Fig. 3.**
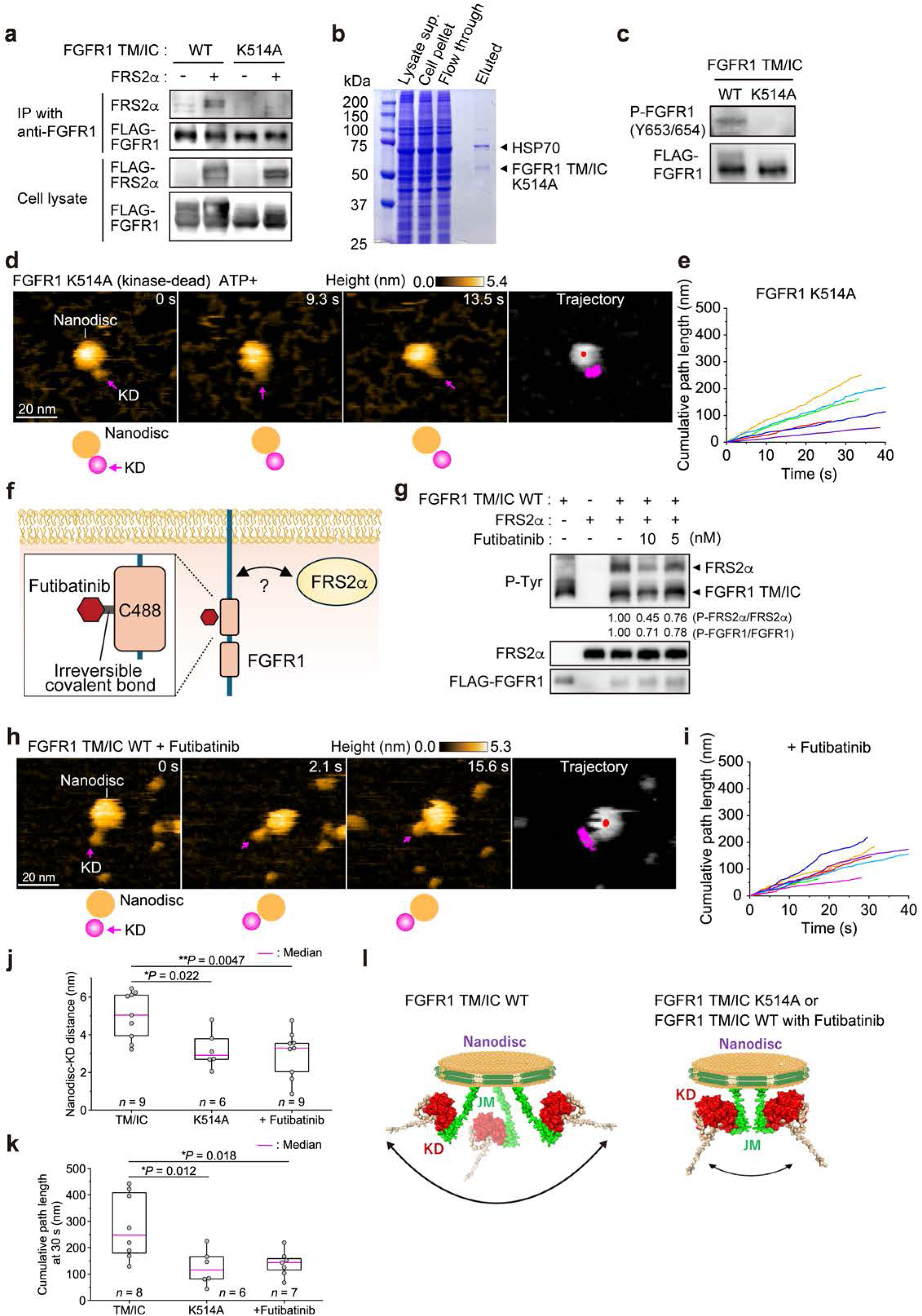
Kinase activity governs receptor conformational dynamics required for productive adaptor engagement. **a,** Co-immunoprecipitation of FLAG-tagged FRS2α with wild-type (WT) or kinase- inactive K514A FGFR1. FRS2α was co-expressed with either FLAG-tagged WT or kinase-inactive FGFR1 (K514A). Immunoprecipitation was performed using antibody against FGFR1, followed by immunoblotting with antibodies against FRS2α or FLAG. **b,** Purified WT and K514A FGFR1 TM/IC are visualized by Coomassie blue staining. **c,** Phosphorylation of purified FGFR1 TM/IC WT and K514A. Immunoblot confirming loss of kinase activity in K514A. **d,** Representative HS-AFM image sequence of membrane- reconstituted K514A FGFR1. The KD remains closely associated with the nanodisc, indicating a compact intracellular configuration. **e,i,** Quantification of the distance between the nanodisc centre and KD. **f,** Schematic illustration of action of FGFR1 inhibitor, futibatinib. **g,** Phosphorylated tyrosine (P-Tyr) was detected after incubation of purified FRS2α (37 ng/μl) and FGFR1 TM/IC WT (10.8 ng/μl) proteins in kinase reaction buffer. Phosphorylation levels of FRS2α and FGFR1 TM/IC, calculated as the band intensity of each phosphorylated protein divided by that of the corresponding total protein, are shown. **h,** Representative HS-AFM image sequences of futibatinib-treated WT FGFR1 showing a compact intracellular configuration resembling K514A. **d,h**, Schematic models corresponding to each image are shown below. The rightmost panels show trajectories of the center of the nanodisc (red point in gray nanodisc) and of the KD (magenta) tracked over approximately 30 s. Scale bars, 20 nm. Height scales are shown above each image series. **j,k**, Quantification of the nanodisc–KD distance (**j**) and cumulative distance (**k**) travelled at 30 s in the indicated conditions. Boxes depict median with upper and lower quartiles and whiskers indicate minimum and maximum. *n* denotes the number of FGFR1 molecules analysed. *P* values were determined by Kruskal-Wallis test with Dunn’s post hoc test. **l,** In FGFR1 WT, the intracellular region undergoes dynamic conformational fluctuations, allowing the KD to move a broad range of positions beneath the plasma membrane. In contrast, loss of kinase activity (K514A) or treatment with a kinase inhibitor restricts the mobility of the JM region and KD.

To determine how kinase activity influences receptor dynamics, K514A FGFR1 was reconstituted into lipid nanodiscs and analysed by HS-AFM. In contrast to the broad conformational dynamics observed for activated wild-type FGFR1, the kinase domain of K514A remained closely associated with the membrane, with minimal separation attributable to the JM region (Fig. 3d and Supplementary Movie 5). Quantitative trajectory analysis demonstrated that the kinase domain explored a markedly smaller conformational space than wild-type FGFR1 (Fig. 3d,e). Consistent with the co-immunoprecipitation results, we did not observe a particle consistent with productive FRS2α engagement under these conditions. Together, these findings indicate that loss of kinase activity constrains the conformational flexibility of the JM region, restricting the dynamic receptor configuration required for productive adaptor engagement.

We next asked whether pharmacological inhibition of FGFR kinase activity produces a similar conformational state. Futibatinib is an irreversible covalent inhibitor of FGFR1–4 that targets Cysteine (C) 488 within the FGFR1 kinase domain and is approved for the treatment of FGFR2 fusion- or rearrangement-positive intrahepatic cholangiocarcinoma^23, 26^ (Fig. 3f). In vitro kinase assays demonstrated that futibatinib inhibited phosphorylation of both FGFR1 and FRS2α in a concentration-dependent manner (Fig. 3g).

HS-AFM imaging revealed that futibatinib-treated FGFR1 adopted a membrane-proximal configuration closely resembling that of the kinase-inactive mutant. The kinase domain remained adjacent to the membrane, the spatial separation provided by the JM region was markedly reduced, and kinase domain motion was substantially restricted (Fig. 3h,i and Supplementary Movie 6). Quantification of the nanodisc-to-kinase-domain distance together with cumulative trajectory analysis confirmed that both K514A and futibatinib significantly reduced the conformational space sampled by FGFR1 (Fig. 3j,k). Under these conditions, we did not observe productive FRS2α engagement. Thus, both genetic and pharmacological suppression of kinase activity converge on the same compact receptor configuration, demonstrating that kinase activity governs the conformational dynamics required for productive adaptor engagement (Fig. 3l).

These findings prompted us to investigate the structural mechanism by which receptor autophosphorylation expands the conformational space of FGFR1 and enables adaptor engagement.

### Autophosphorylation releases JM autoinhibition to enable productive adaptor engagement

Having established that kinase activity governs receptor conformational dynamics, we next sought to define the structural mechanism underlying this dynamic regulation. To this end, we performed all-atom molecular dynamics (MD) simulations of membrane-embedded FGFR1 TM/IC in three conformational states: wild-type (WT), the kinase-inactive mutant (K514A), and a phosphomimetic mutant (Y5D), in which the five major autophosphorylation sites within the kinase domain were replaced by aspartic acid (D)^25^ (Fig. 4a–c). Simulations were initiated from AlphaFold3-predicted structures embedded in a lipid bilayer.

**Fig. 4.**
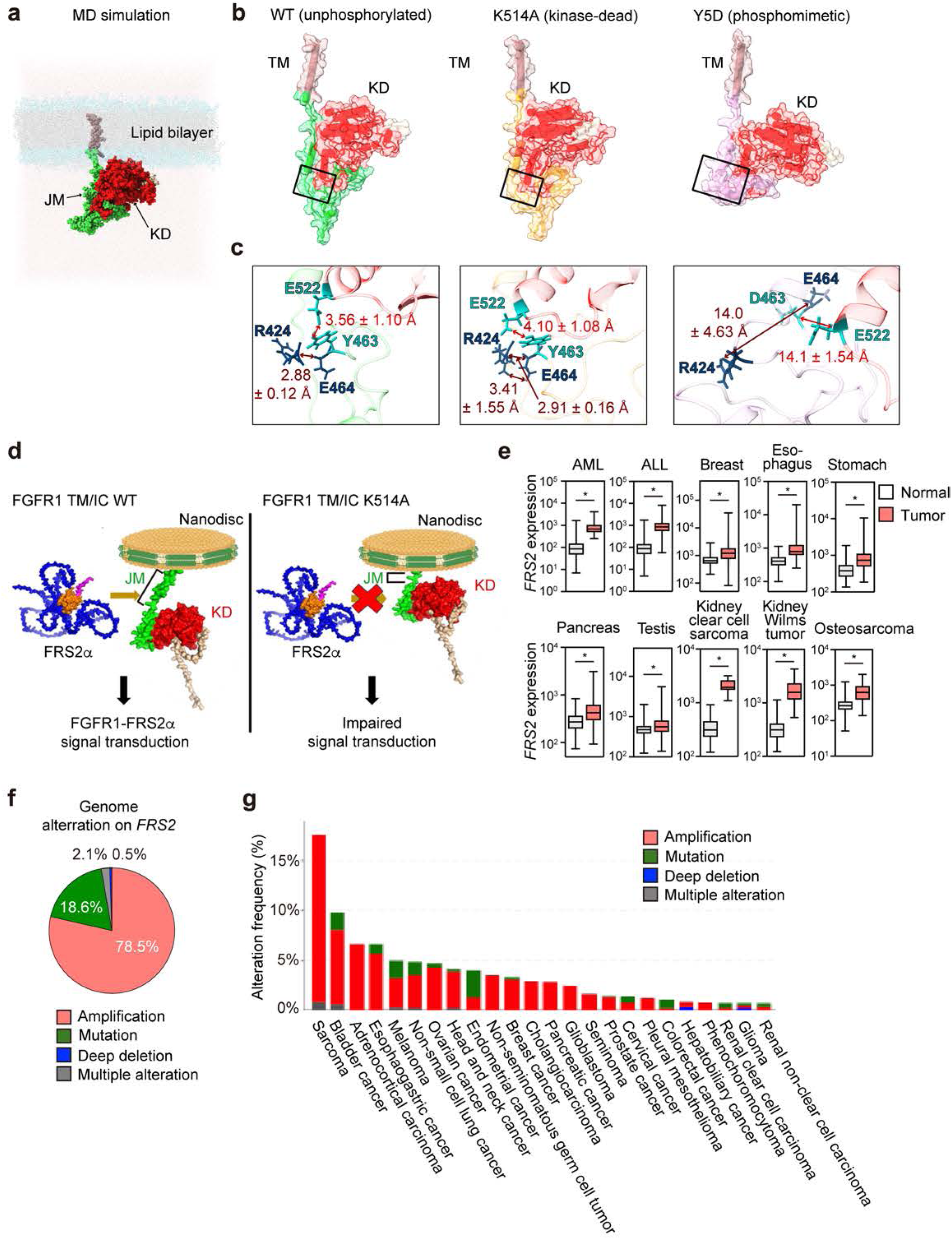
Autophosphorylation relieves juxtamembrane autoinhibition to enable productive adaptor engagement. **a,** Molecular dynamics (MD) simulation of membrane-embedded FGFR1 TM/IC in the WT, kinase-inactive K514A and phosphomimetic Y5D states. **b,c,** Representative MD simulation snapshots (**b**) and corresponding structural interactions (**c**) showing that WT and K514A maintain intramolecular contacts between the JM region and KD, whereas these interactions are disrupted in the phosphomimetic Y5D mutant (Tyr463, Tyr583, Tyr585, Tyr653 and Tyr654 replaced by Asp), all comprising residues 377–773, extracted from the final snapshot of a representative 200-ns trajectory. The KD is red. Distances represent the mean ± s.d. of three independent trajectories. The backbone hydrogen bond between Arg424 and Glu464, and the contact between Tyr463 and Glu522, present in WT and K514A, are lost in Y5D. The WT protein was simulated in its unphosphorylated state and thus corresponds to the kinase-inactive condition, whereas Y5D mimics the autophosphorylated WT protein. Structures were rendered with UCSF ChimeraX (v. 1.11.1). **d,** Proposed model for kinase-dependent regulation of FGFR1 signaling. In the inactive state, the JM region adopts a conformationally restricted configuration that limits productive adaptor engagement. Autophosphorylation releases this structural restraint, allowing the receptor to sample an expanded conformational space that enables productive engagement of FRS2α. Genetic (K514A) or pharmacological (futibatinib) inhibition maintains the compact receptor configuration. **e,** Expression of the gene encoding FRS2α (*FRS2*) in normal tissues and human cancers. Boxes depict median with upper and lower quartiles and whiskers indicate minimum and maximum. *P* values were determined by two-tailed Mann-Whitney test. **f,g,** Frequency (**f**) and distribution (**g**) of *FRS2* genomic alterations across human cancers. Gene amplification is the predominant mechanism of *FRS2* alteration.

In both the unphosphorylated WT and K514A receptors, the JM region remained closely associated with the kinase domain throughout the simulations (Fig. 4b,c). The backbone amide of Arginine (R) 424 formed a hydrogen bond with the backbone carbonyl of Glutamic acid (E) 464, with a donor–acceptor (N···O) distance of 2.88 ± 0.12 Å (WT) and 3.41 ± 1.55 Å (K514A) (Fig. 4c), and Tyrsine (Y) 463 was positioned close to E522 (3.56 ± 1.10 Å in WT and 4.10 ± 1.08 Å in K514A). Together, these contacts held the JM region against the KD, keeping the intracellular module in a compact configuration. By contrast, in the phosphomimetic Y5D mutant, both contacts were disrupted: the R424–E464 distance increased to 14.0 ± 4.63 Å and the D463–E522 distance to 14.1 ± 1.54 Å (Fig. 4c). The electrostatic repulsion between the negatively charged D463 and E522 further separated the JM region from the KD, releasing the intracellular module from its compact state. These structural rearrangements provide a mechanistic explanation for the HS-AFM observations, indicating that receptor autophosphorylation relieves JM-mediated autoinhibition and permits the kinase domain to explore a substantially broader conformational space.

Based on the HS-AFM, biochemical and MD analyses, we propose that kinase activity regulates productive adaptor engagement by controlling the conformational flexibility of the intrinsically disordered JM region (Fig. 4d). In the absence of kinase activity, the JM region remains constrained against the kinase domain, limiting productive engagement of FRS2α. Autophosphorylation releases this structural restraint, allowing dynamic opening of the receptor and productive adaptor engagement.

To assess the broader biological relevance of this mechanism, we next examined the status of FRS2α in human cancer. Analysis of public transcriptomic datasets demonstrated that the gene encoding the adaptor protein FRS2α (*FRS2*) is significantly overexpressed in multiple tumour types compared with corresponding normal tissues ^27^ (Fig. 4e). Consistent with this observation, analysis of cancer genomics datasets identified gene amplification as the predominant mechanism of *FRS2* alteration across diverse human malignancies ^28,29,30^ (Fig. 4f,g). These observations indicate that the principal adaptor regulated by kinase-dependent receptor dynamics is frequently dysregulated in human cancer, highlighting the biological and clinical relevance of the signaling mechanism identified here.

Collectively, our findings support a model in which receptor autophosphorylation releases a previously constrained conformation of the intrinsically disordered FGFR1 JM region, enabling the conformational dynamics required for productive FRS2α engagement. Conversely, genetic or pharmacological inhibition of kinase activity maintains the receptor in a compact, conformationally restricted state that is incompatible with efficient receptor-to-adaptor signaling.

## Discussion

Signal transduction is initiated through a series of molecular events that occur within seconds after receptor activation^13^, yet the earliest physical step linking receptor activation to intracellular signaling has remained largely inaccessible to direct observation. Structural biology has defined the molecular architecture of receptor tyrosine kinases, whereas biochemical studies have established the signaling pathways that follow receptor activation. By directly visualizing individual FGFR1 molecules in real time, our study reveals a previously unseen layer of receptor signaling: kinase-dependent conformational dynamics of an intrinsically disordered receptor region that enable productive adaptor engagement. Rather than functioning as a passive flexible linker, the JM region actively explores the intracellular space following receptor autophosphorylation, thereby allowing efficient engagement of the signaling adaptor FRS2α. We therefore propose that regulated conformational dynamics constitute the physical mechanism that couples receptor activation to the initiation of intracellular signal transduction.

Previous structural studies demonstrated that JM and C-terminal regions of several RTKs interact with the kinase domain to stabilize inactive conformations. Similar autoinhibitory interactions have been described for EGFR, EphB2 and FGFR family members ^30,31,32,33,34^. Our molecular dynamics simulations provide direct structural support for this model by showing that autophosphorylation disrupts intramolecular interactions that constrain the JM region, thereby allowing it to sample a substantially broader conformational space. Together with the HS-AFM observations, these findings support a model in which receptor autophosphorylation relieves JM autoinhibition to generate the dynamic receptor configuration required for productive adaptor engagement.

Although this study focuses on the FGFR1–FRS2α signaling axis, the underlying mechanism is likely to extend beyond this receptor system. Intrinsically disordered intracellular regions are a common feature of many RTK and frequently serve as docking platforms for signaling proteins. FRS2α itself also mediates signaling downstream of additional receptors, including neurotrophin receptors and RET ^17^. Likewise, the C-terminal intrinsically disordered region of FGFR1 recruits signaling molecules such as PLCγ and STAT1^15^, whereas the disordered C-terminal region of EGFR participates in multivalent interactions with Grb2 that promote higher-order signaling assemblies and phase separation. Whether these diverse signaling events are similarly regulated by kinase-dependent conformational dynamics will be an important question for future investigation^8, 35^. HS-AFM now provides an opportunity to directly visualize these dynamic signaling processes at the single-molecule level.

One limitation of the present study is that FRS2α was immobilized on the mica surface to enable high-resolution imaging. Consequently, although HS-AFM directly visualized productive adaptor engagement, the current experimental configuration does not permit quantitative analysis of association frequency, dwell time or dissociation kinetics. Likewise, receptor dimerization was not analysed under the present conditions. Future improvements in the HS-AFM substrate should enable visualization of increasingly complex receptor signaling assemblies, including dynamic interactions among receptors, adaptors and downstream signaling proteins.

Structure-guided development of tyrosine kinase inhibitors has transformed the treatment of many human cancers^6^. However, most currently available inhibitors target the catalytic kinase domain and frequently lose efficacy because of acquired resistance mutations. Our findings raise the possibility of an alternative therapeutic strategy based on modulating receptor conformational dynamics rather than catalytic activity itself. Compounds that alter the dynamic behaviour of intrinsically disordered receptor regions could potentially disrupt productive adaptor engagement while remaining mechanistically distinct from conventional inhibitors that binds to the kinase domain. More broadly, our work establishes HS-AFM as a platform for directly visualizing receptor signaling in real time and provides a framework for discovering compounds that regulate signal transduction by reshaping protein conformational dynamics.

By revealing the dynamic molecular events that physically couple receptor activation to adaptor engagement, our study establishes a new framework for understanding receptor tyrosine kinase signaling and opens new opportunities for visualizing and therapeutically targeting dynamic mechanisms of signal transduction.

## Material and Method

### Cell culture and transfection

HEK293T cells (ATCC) were cultured in Dulbecco’s Modified Eagle Medium (DMEM; Nacalai Tesque, Kyoto, Japan) supplemented with 10% fetal bovine serum (FBS; Thermo Fisher Scientific). Expi293F cells (Thermo Fisher Scientific) were maintained in Expi293 Expression Medium (Thermo Fisher Scientific) at 37°C in a humidified incubator with 8% CO₂ and shaking at 125 rpm. Plasmid transfection of Expi293F cells was performed using the ExpiFectamine™ 293 Transfection Kit (Thermo Fisher Scientific) according to the manufacturer’s instructions. Eighteen to 20 h after transfection, ExpiFectamine™ 293 Transfection Enhancer 1 and ExpiFectamine™ 293 Transfection Enhancer 2 (Thermo Fisher Scientific) were added to the cultures. Cells were harvested 5 days after transfection, and cell pellets were flash-frozen in liquid nitrogen and stored at −80°C until recombinant protein purification.

### Plasmids

A truncated human FGFR1 construct lacking the extracellular domain (FGFR1 TM/IC- WT; amino acids 377–822) was cloned into the mammalian expression vector pCAGGS- His7-FLAG using the NheI and EcoRV restriction sites. The human FGFR1 TM/IC-K514A fragment (amino acids 377–822) was synthesized as a GeneArt String (Thermo Fisher Scientific). XbaI and EcoRV restriction sites were introduced at the 5′ and 3′ ends, respectively, by polymerase chain reaction (PCR), and the fragment was subsequently cloned into the NheI and EcoRV sites of the pCAGGS-His7-FLAG vector. The human FRS2α fragment (amino acids 1–508) containing the G2A mutation, which improves protein solubility during purification, was synthesized as a GeneArt String (Thermo Fisher Scientific). A NheI site at the 5′ end and a PvuII site at the 3′ end were introduced by PCR, and the fragment was cloned into the NheI and EcoRV sites of the pCAGGS- His7-FLAG vector. *Escherichia coli* DH5α cells (Toyobo, Osaka, Japan) were used as the host strain for molecular cloning.

### Purification of recombinant proteins

Cell pellets of Expi293F cells expressing FGFR1 TM/IC-WT, FGFR1 TM/IC-K514A, or FRS2α-G2A were resuspended in a lysis buffer containing 50 mM Tris-HCl (pH 7.5), 150 mM NaCl, 1 mM EGTA, and 1× Protease Inhibitor Cocktail (Nacalai Tesque). The cells were lysed using a Q125 micro ultrasonic homogenizer (Waken B Tech), followed by centrifugation to separate the soluble and insoluble fractions. The supernatant was collected and incubated with Anti-FLAG M2 Affinity Gel (Millipore) at 4°C for 1 h with gentle rotary agitation. After centrifugation, the resin was washed five times with Tris-buffered saline (TBS). FLAG-tagged proteins were eluted by incubating the resin with 100 μg/mL FLAG peptide (Millipore). The purified proteins were used for mass spectrometry, in vitro kinase assays, and high-speed atomic force microscopy (HS-AFM) observations.

### Mass spectrometry

The eluates obtained by purification with Anti-FLAG M2 Affinity Gel (Millipore) were subjected to mass spectrometry analysis. Proteins in the eluates were separated by sodium dodecyl sulfate–polyacrylamide gel electrophoresis (SDS-PAGE), and the resulting protein bands were excised from the gel.

Excised gel pieces were destained as appropriate and washed twice with 50% acetonitrile in 25 mM ammonium bicarbonate. Proteins in the gel were reduced with 50 mM TCEP in 25 mM ammonium bicarbonate at 60 °C for 10 min and subsequently alkylated with 500 mM iodoacetamide at room temperature for 1 h in the dark. After washing and dehydration with acetonitrile, the samples were rehydrated with a trypsin solution containing 100 ng of trypsin and incubated overnight at 30 °C. The resulting peptide solution was collected, and residual peptides were extracted with 1% TFA. The combined extracts were desalted using Pierce C18 Spin Tips (#84850; ThermoFisher Scientific), concentrated by vacuum centrifugation, reconstituted in 5% acetonitrile containing 0.1% TFA, and filtered through an Ultrafree-MC centrifugal filter (UFC30SV00; Merck) before LC–MS/MS analysis.

The trypsin-digested peptides were analyzed by Orbitrap QE plus (ThermoFisher Scientific) with nano-liquid chromatography (EASY-nLC 1200; ThermoFisher Scientific). The purified peptides were loaded and separated on the Aurora column (25 cm x 75 µm ID, 1.6 mm C18; Ionoptics) with a linear acetonitrile gradient (0-40%) in 0.1% formic acid for 60 min at a flow rate of 300 nL min-1. The peptide ions were detected by Orbitrap QE plus MS; ThermoFisher Scientific) in the data-dependent acquisition mode with the installed Xcalibur software (ThermoFisher Scientific). Full-scan mass spectra were acquired in the MS over 375-1,500 m/z with resolution of 70,000. The MS/MS searches were carried out using SEQUEST HT search algorithms against the Homo sapiens (Swiss prot. Tax ID 9609) protein database using Proteome Discoverer (PD) (Version 3.0; Thermo Fisher Scientific).

### *In vitro* kinase assay

Kinase assays were performed in a reaction buffer containing 40 mM Tris-HCl (pH 7.5), 20 mM MgCl₂, 50 μM dithiothreitol (DTT), and 50 μM adenosine triphosphate (ATP; Promega). Purified FGFR1 TM/IC and FRS2α proteins were mixed in the reaction buffer with or without futibatinib (1-10 nM, Selleck) and incubated at 37°C for 1 h. Reactions were terminated by the addition of SDS sample buffer, and protein phosphorylation was analyzed by western blotting.

### Lipid reconstitution of FGFR1

Wild-type and K514A FGFR1 TM/IC were reconstituted into lipid bilayers by a modified nanodisc assembly protocol based on the manufacturer’s protocol (Sigma‒Aldrich, USA), with adjustments as previously described^21^. Soy-derived phospholipids (40 μg, #11145, Sigma‒Aldrich) were dissolved in chloroform, dried under N_2_, and resuspended in 50 μl of buffer containing 20 mM HEPES-NaOH (pH 7.4), 100 mM NaCl, and 4% n-dodecyl-β-D-maltoside (DDM); the suspension was sonicated (1 min, probe sonicator, TAITEC, Japan) to homogeneity. Solubilized FGFR1 (100 ng/μl, 30 μl) was combined with membrane scaffold protein MSP1E3D1 (Sigma‒Aldrich; final concentration, 1 mg/ml) and incubated for 1 hour at 4°C with rotation. Detergent removal and bilayer formation were initiated with Bio-Beads SM-2 (30 mg; #1523920, Bio-Rad) and allowed to proceed overnight at 4°C. Size-exclusion chromatography, typically used to isolate ∼15-nm nanodiscs, was omitted to optimize samples for HS-AFM; this yielded flat membrane structures under 15 nm in diameter, suitable for high-resolution imaging of the receptor.

### HS-AFM observations

Topographic imaging was performed using a custom-built HS-AFM operated in tapping mode^21^. The instrument incorporated an optical beam deflection (OBD) system equipped with a 780-nm infrared laser (0.7 mW), focused through a 60× objective lens (CFI S Plan Fluor ELWD, Nikon, Japan) onto the gold-coated backside of a cantilever (AC7, MNOIC, Japan); the reflected signal was captured by a segmented PIN photodiode. Cantilevers had dimensions of 6–7 µm (length) × 2 µm (width) × 100 nm (thickness), a spring constant of 0.1 N/m, a resonant frequency of ∼500 kHz in liquid, and a quality factor of ∼2. To ensure high-resolution imaging, an amorphous carbon tip (∼250 nm in length, <1 nm apex radius) was fabricated onto each cantilever by electron beam deposition (EBD). During observations, the free oscillation amplitude was maintained at ∼1 nm, and the set-point amplitude was tuned to ∼90% of the free amplitude for stable feedback control. APTES-functionalized mica (AP-mica) served as the substrate, prepared by incubating freshly cleaved mica with 0.005% (v/v) 3- aminopropyltriethoxysilane (APTES, #440140, Sigma-Aldrich) in Milli-Q water for 3 min at room temperature. The substrate was then rinsed with 40 μl of Milli-Q water followed by the imaging buffer (50 mM Tris-HCl [pH 7.4], 150 mM NaCl). AP-mica was used to immobilize the negatively charged nanodiscs through electrostatic interactions.

All HS-AFM imaging was performed in imaging buffer consisting of 50 mM Tris-HCl (pH 7.4) and 150 mM NaCl at 24–26 °C. Unless otherwise stated, samples were deposited onto a freshly prepared AP-mica substrate before imaging.

*Nanodiscs containing FGFR1 TM/IC*. Nanodiscs reconstituted with FGFR1 TM/IC (WT or K514A) were incubated at 37 °C for 1 h in kinase assay buffer containing 50 μM ATP, followed by incubation at 4 °C overnight.

*FRS2α.* Purified FRS2α was diluted to 20 nM in 50 mM Tris-HCl (pH 7.4) and 150 mM NaCl and deposited onto freshly cleaved bare mica.

*FGFR1–FRS2α complex.* FGFR1 TM/IC WT reconstituted into nanodiscs was mixed with purified FRS2α at a 1:1 molar ratio and incubated in kinase assay buffer containing 50 μM ATP at 37 °C for 1 h.

*FGFR1 with futibatinib.* FGFR1 TM/IC WT reconstituted into nanodiscs was preincubated with 10 nM futibatinib in ATP-free kinase assay buffer for 15 min at room temperature. ATP was then added to a final concentration of 50 μM, and the mixture was incubated at 37 °C for 1 h, followed by further incubation at 4 °C overnight.

Images in Fig. 1 **i** and **j** were recorded at a frame rate of 3.3 frames/s in buffer containing (50 mM Tris-HCl [pH 7.4], 150 mM NaCl) at room temperature (24–26°C). Scan area, 80 × 64 nm^2^ (100 × 80 pixels).

Images in Fig. 2 **e** and **f** were recorded at a frame rate of 3.3 frames/s in buffer containing 50 mM Tris-HCl (pH 7.4), 150 mM NaCl at room temperature (24–26°C). Scan area, 80 × 64 nm^2^ (100 × 80 pixels).

Images in Fig. 3 **d** and **h** were recorded at a frame rate of 3.3 frames/s in buffer containing 50 mM Tris-HCl (pH 7.4), 150 mM NaCl at room temperature (24–26°C). Scan area, 80 × 64 nm^2^ (100 × 80 pixels).

### HS-AFM image processing and data analysis

We processed all HS-AFM images using the Fiji distribution of ImageJ^36^. To enhance the signal-to-noise ratio, a mean filter with a 0.5-pixel radius was applied to each frame, and background fluctuations were removed using the “Subtract Background” plugin with a rolling-ball radius of 200 pixels. Lateral drift between successive frames was corrected using the Template Matching and Slice Alignment plugin^37^. Centroids of the nanodiscs and kinase domains were tracked semi-manually across all experimental conditions using the MTrackJ plugin^38^. The distance between the kinase domain and the nanodiscs was defined as the displacement between their centroids minus the nanodisc radius; for each molecule, this distance was calculated over ∼100 successive frames, and the median value was used to represent individual molecular behavior. Analysis was performed on the following datasets: wild-type FGFR1 TM/IC (*n* = 9 molecules, 955 total frames; average 106, range 62-151 frames per molecule), the K514A mutant (*n* = 6 molecules, 765 total frames; average 128, range 89-176 frames per molecule), and wild-type FGFR1 TM/IC treated with futibatinib (*n* = 9 molecules, 938 total frames; average 104, range 44-210 frames per molecule). Kinase domain flexibility was assessed using the Len [nm] parameter from MTrackJ, representing the accumulated path length of the tracked centroids; total Len values at t = 30 s were compared across conditions to determine statistical significance.

### In silico analysis of protein structures

Three-dimensional structures of FGFR1 TM/IC and FRS2α were predicted using AlphaFold3 (v3.0.0)^24^. For FGFR1 TM/IC (residues 377–822), structural models of the WT, K514A, and phosphomimetic Y5D mutant (Y463D/Y583D/Y585D/Y653D/Y654D) were generated. For each construct, the model with the highest ranking score was selected for subsequent analyses. Intrinsically disordered regions (IDRs) were predicted using the PrDOS Protein Disorder Prediction System (disorder probability > 0.5) ^39^. Predicted structures were visualized using PyMOL Molecular Graphics System (Version 3.10.17, Schrödinger, LLC) or UCSF ChimeraX (v1.11.1) ^40^.

### Molecular dynamics (MD) simulation of FGFR1

To compare the structural stability and mutation-induced conformational changes of the FGFR1 TM/IC region among the wild-type (WT), K514A and Y5D constructs predicted by AlphaFold3, we performed all-atom MD simulations. Because no reliable structure could be predicted for the C-terminal linker (residues 774–822), this region was excluded, and residues 377–773 were used as the initial structure. Simulations were run with GROMACS 2024.2^41^. Systems were built with CHARMM-GUI Membrane Builder^42^, with the position and orientation of each protein in the bilayer set by automated placement using the integrated PPM server integrated into CHARMM-GUI^43^. Each protein was embedded in a pure 1-palmitoyl-2-oleoyl-sn-glycero-3-phosphocholine (POPC) bilayer that was symmetric in both leaflets. Protonation states of titratable residues were assigned at neutral pH using the default CHARMM-GUI settings. Each system was solvated with TIP3P water, and 0.15 M Na^+^ and Cl^-^ ions were added to neutralize the net charge. The AMBER ff19SB force field^44^ was used for the protein, Lipid21^45^ for POPC, and the TIP3P model ^46^for water, and the Joung–Cheatham parameters for monovalent ions^47^ for Na⁺ and Cl⁻; force field parameters and simulation inputs were generated with the CHARMM-GUI FF-Converter^48^. Three-dimensional periodic boundary conditions were applied. The system comprised one FGFR1 TM/IC domain, 741 POPC molecules, 102,843 water molecules, 282 Na+ ions, and 284 Cl− ions. The histidine residues were assumed to be in the protonated state at pH = 7, and two cysteine residues were assumed to have a disulfide bond when they were close together. The total number of atoms in the system was 414,683. Simulation box dimensions are 160.081 × 160.081 × 171.434 Å^3^.

Energy minimization was performed using the steepest-descent algorithm for a maximum of 10,000 steps or until the maximum force fell below 1,000 kJ mol⁻¹ nm⁻¹. During minimization, short-range electrostatic and van der Waals interactions were truncated at 1.6 nm. Long-range electrostatic interactions were treated with the particle mesh Ewald method^49^. Positional restraints were applied to the protein backbone, protein side chains, and lipids, together with dihedral restraints on the protein.

The systems were subsequently equilibrated at 300 K in seven successive stages totalling 125 ps. The first two stages consisted of 5-ps NVT equilibration runs, followed by four semi-isotropic NPT stages at 1 bar lasting 5, 10, 20, and 40 ps, respectively, in which the simulation box dimensions in the membrane plane and along the membrane-normal axis were scaled independently. A final 40-ps NVT equilibration stage was then performed. The temperature was maintained at 300 K using the velocity-rescaling thermostat^50^ with a coupling time constant of 1.0 ps, with the protein, the lipid bilayer, and the solvent (water and ions) treated as separate temperature-coupling groups. The pressure was maintained at 1 bar using semi-isotropic coupling with the C-rescale barostat^51^, with a coupling time constant of 5.0 ps and an isothermal compressibility of\ 4.5 × 10⁻⁵ bar⁻¹. A time step of 1 fs was used during the first three stages and 2 fs thereafter. Initial velocities were assigned from a Maxwell–Boltzmann distribution at 300 K at the beginning of the first NVT stage. Positional and dihedral restraints were progressively reduced over the seven equilibration stages. Positional force constants were decreased from 4,000 to 50 kJ mol⁻¹ nm⁻² for the protein backbone, from 2,000 to 0 kJ mol⁻¹ nm⁻² for the protein side chains, and from 1,000 to 0 kJ mol⁻¹ nm⁻² for the lipids. Protein dihedral restraints were likewise reduced stepwise (**Supplementary table**).

After equilibration, all restraints were removed, and 200-ns simulations were performed in the NPT ensemble at 300 K and 1 bar with a time step of 2 fs using the same thermostat and barostat settings as described above. Long-range electrostatic interactions were treated with the particle mesh Ewald method, using a real-space cutoff of 1.0 nm. A cutoff of 1.0 nm was applied to van der Waals interactions, and a long-range dispersion correction was applied to both the energy and the pressure. Neighbor lists were generated with the Verlet cutoff scheme. The list was updated every 0.160 ps. All bonds involving hydrogen atoms were constrained with the LINCS algorithm^52^, and water geometry was constrained with SETTLE ^53^.

For each of the three variants (WT, K514A, and 5YD), three independent 200-ns simulations were performed using different random seeds for initial velocity assignment, yielding nine trajectories and an aggregate simulation time of 1.8 μs. Coordinates and energies were recorded every 1 ns and 20 ps, respectively. Of the 200 ns in each independent simulation, the first 100 ns were discarded as equilibration calculations, and the latter 100 ns were used for structural analysis.

### Immunoprecipitation

HEK293T cells were transfected with expression vectors encoding FRS2α-G2A and FGFR1 TM/IC (WT or K514A) using Lipofectamine 2000 (Thermo Fisher Scientific) and cultured for 2 days to allow protein expression. Cells were lysed in a lysis buffer containing 10 mM Tris-HCl (pH 7.5), 5 mM EDTA, 150 mM NaCl, 1% Triton X-100, 1× Protease Inhibitor Cocktail (Nacalai Tesque), and 1× Phosphatase Inhibitor Cocktail (Nacalai Tesque), followed by incubation at 4°C for 20 min with rotation. Cell lysates were centrifuged, and the resulting supernatants were incubated with 1 μg of anti-FGFR1 polyclonal antibody (#30358-1-AP, Proteintech) for 90 min at 4°C with rotation. Protein G Sepharose 4 Fast Flow (GE Healthcare) was then added, and the mixtures were incubated for an additional 90 min at 4°C with rotation. The Sepharose beads were washed five times with lysis buffer, and the immunoprecipitated proteins were eluted by the addition of SDS sample buffer. The eluted proteins were subsequently analyzed by western blotting.

### Western blot

Western blotting was performed using primary antibodies against the FLAG tag (clone M2, 1:2,000, MilliporeSigma), FRS2α (1:500, #MAB4069, R&D Systems), phospho- FRS2α (Tyr196) (1:1,000, #3864, Cell Signaling Technology), phospho-FGFR1 (Tyr653/654) (1:1,000, #3471, Cell Signaling Technology), and phosphotyrosine (PY20, 1:4,000, BD Transduction Laboratories), followed by horseradish peroxidase-conjugated secondary antibodies (Cytiva). Protein signals were visualized using Immobilon Western Chemiluminescent HRP Substrate (Millipore) and detected with an ImageQuant LAS 4000 imaging system (Cytiva).

### Analysis of gene expression and genomic alterations of FRS2α

FRS2α expression in normal and tumor tissues across different cancer types was analyzed using TNMplot V2^27^. Genomic alterations of FRS2 were analyzed using cBioPortal^28–30^ based on The Cancer Genome Atlas (TCGA) PanCancer Atlas studies, including adrenocortical carcinoma, cholangiocarcinoma, bladder urothelial carcinoma, colorectal adenocarcinoma, breast invasive carcinoma, brain lower grade glioma, glioblastoma multiforme, cervical squamous cell carcinoma, esophageal adenocarcinoma, stomach adenocarcinoma, uveal melanoma, head and neck squamous cell carcinoma, kidney renal clear cell carcinoma, kidney chromophobe, kidney renal papillary cell carcinoma, liver hepatocellular carcinoma, lung adenocarcinoma, lung squamous cell carcinoma, diffuse large B-cell lymphoma, acute myeloid leukemia, ovarian serous cystadenocarcinoma, pancreatic adenocarcinoma, mesothelioma, prostate adenocarcinoma, skin cutaneous melanoma, pheochromocytoma and paraganglioma, sarcoma, testicular germ cell tumors, thymoma, thyroid carcinoma, uterine corpus endometrial carcinoma, and uterine carcinosarcoma.

### Quantification and statistical analysis

Statistical analyses were performed using GraphPad Prism and Igor Pro 10 (WaveMetrics, USA). For all statistical tests, the significance level was set at α = 0.05. Data normality and homogeneity of variance were assessed using the Shapiro–Wilk test and Bartlett’s test, respectively. Comparisons among three experimental groups were performed using the Kruskal–Wallis test followed by Dunn’s multiple-comparison test. Statistical significance was defined as follows: \**P* < 0.05, \*\**P* < 0.01, and \*\*\**P* < 0.001; *P* > 0.05 was considered not significant (NS). Detailed information on the statistical tests and sample sizes for each experiment is provided in the corresponding figure legends.

### Use of large language models

Chat-GPT and Claude were employed for proofreading the text.

## Acknowledgements

We thank Y. Mikami, Y. Okada, and Y. Kamide (Kanazawa University) for their essential administrative and laboratory support. We thank Dr. T. Nishiuchi and members in Bioscience Core Facility, Research Center of Experimental Modeling of Human Disease, Kanazawa University, for their technical assistance on mass spectrometry. This research was supported by the World Premier International Research Center Initiative (WPI), MEXT, Japan (to M.S.), JSPS KAKENHI grant numbers JP25H00972 (to M.S.), JP22H04926 Advanced Bioimaging Support (ABiS) (to M.S.), the Mochida Memorial Foundation for Medical and Pharmaceutical Research (to M.S.), the Uehara Memorial Foundation (to M.S.), the Naito Foundation (to M.S.), the Takeda Science Foundation, JST SPRING (JPMJSP2135 to K.S.), and JST ERATO (JPMJER2403 to M.S.). MD simulations were partly conducted on a supercomputer at the Research Center for Computational Science in Okazaki, Japan (project ID: 26-IMS-C167). This work was also supported by JSPS KAKENHI (JP23K18237, JP24K02304 and JP25K22575 to N.G.); the AMED Project for Cancer Research and Therapeutic Evolution (21447913, 21446781, 24028177 and 24021759 to N.G.); the Uehara Memorial Foundation (to N.G.); the Princess Takamatsu Cancer Research Fund (to N.G.); the Takeda Science Foundation (to N.G.); the Chugai Foundation for Life Science (to N.G.); and the Yasuda Medical Foundation (to N.G.). This work was supported in part by the MEXT Promotion of Development of a Joint Usage/Research System Project, Coalition of Universities for Research Excellence Program (CURE; JPMXP1323015484 to T.H. and N.G.).

## Author contributions

M.S. and N.G. conceived the project. Y.M., H.Z., T.H., Sarenqiqige, M.S. and N.G. performed AFM imaging and interpreted the data. Y.M., H.Z., T.H., Sarenqiqige and N.G. performed biochemical experiments and interpreted the data. K.S., Y.N. and K.M. established and provided the mammalian expression system. K.S., T.S. and M.S. performed molecular dynamics simulations and interpreted the data. Y.M., H.Z., T.H., K.S., T.S., M.S. and N.G. wrote the manuscript.

## Competing interests

The authors declare no competing interests.

---

**Supplementary table.** Information on the Equilibration System for MD Simulations.

|  | Step 1 | Step 2 | Step 3 | Step 4 | Step 5 | Step 6 | Step 7 |
| --- | --- | --- | --- | --- | --- | --- | --- |
| Ensemble | NVT | NVT | NPT | NPT | NPT | NPT | NVT |
| Time step | 1 fs | 1 fs | 1 fs | 2 fs | 2 fs | 2 fs | 2 fs |
| Number of steps | 5000 | 5000 | 5000 | 5000 | 10000 | 20000 | 20000 |
| Temperature control | V-rescale | V-rescale | V-rescale | V-rescale | V-rescale | V-rescale | V-rescale |
| Pressure control | None | None | C-rescale | C-rescale | C-rescale | C-rescale | None |
| Backbone constraints | 4,000 kJ mol<sup>-1</sup> nm<sup>-2</sup> | 2,000 kJ mol<sup>-1</sup> nm<sup>-2</sup> | 1,000 kJ mol<sup>-1</sup> nm<sup>-2</sup> | 500 kJ mol<sup>-1</sup> nm<sup>-2</sup> | 200 kJ mol<sup>-1</sup> nm<sup>-2</sup> | 50 kJ mol<sup>-1</sup> nm<sup>-2</sup> | 50 kJ mol<sup>-1</sup> nm<sup>-2</sup> |
| Side chain constraints | 2,000 kJ mol<sup>-1</sup> nm<sup>-2</sup> | 1,000 kJ mol<sup>-1</sup> nm<sup>-2</sup> | 500 kJ mol<sup>-1</sup> nm<sup>-2</sup> | 200 kJ mol<sup>-1</sup> nm<sup>-2</sup> | 50 kJ mol<sup>-1</sup> nm<sup>-2</sup> | 0 | 0 |
| Lipid constraints | 1,000 kJ mol<sup>-1</sup> nm<sup>-2</sup> | 400 kJ mol<sup>-1</sup> nm<sup>-2</sup> | 400 kJ mol<sup>-1</sup> nm<sup>-2</sup> | 200 kJ mol<sup>-1</sup> nm<sup>-2</sup> | 40 kJ mol<sup>-1</sup> nm<sup>-2</sup> | 0 | 0 |
| Bipolar constraints | 1,000 kJ mol<sup>-1</sup> rad<sup>-2</sup> | 400 kJ mol<sup>-1</sup> rad<sup>-2</sup> | 200 kJ mol<sup>-1</sup> rad<sup>-2</sup> | 200 kJ mol<sup>-1</sup> rad<sup>-2</sup> | 100 kJ mol<sup>-1</sup> rad<sup>-2</sup> | 0 | 0 |
| Bond length constraints | LINCS | LINCS | LINCS | LINCS | LINCS | LINCS | LINCS |

**Extended Data Fig.1.**
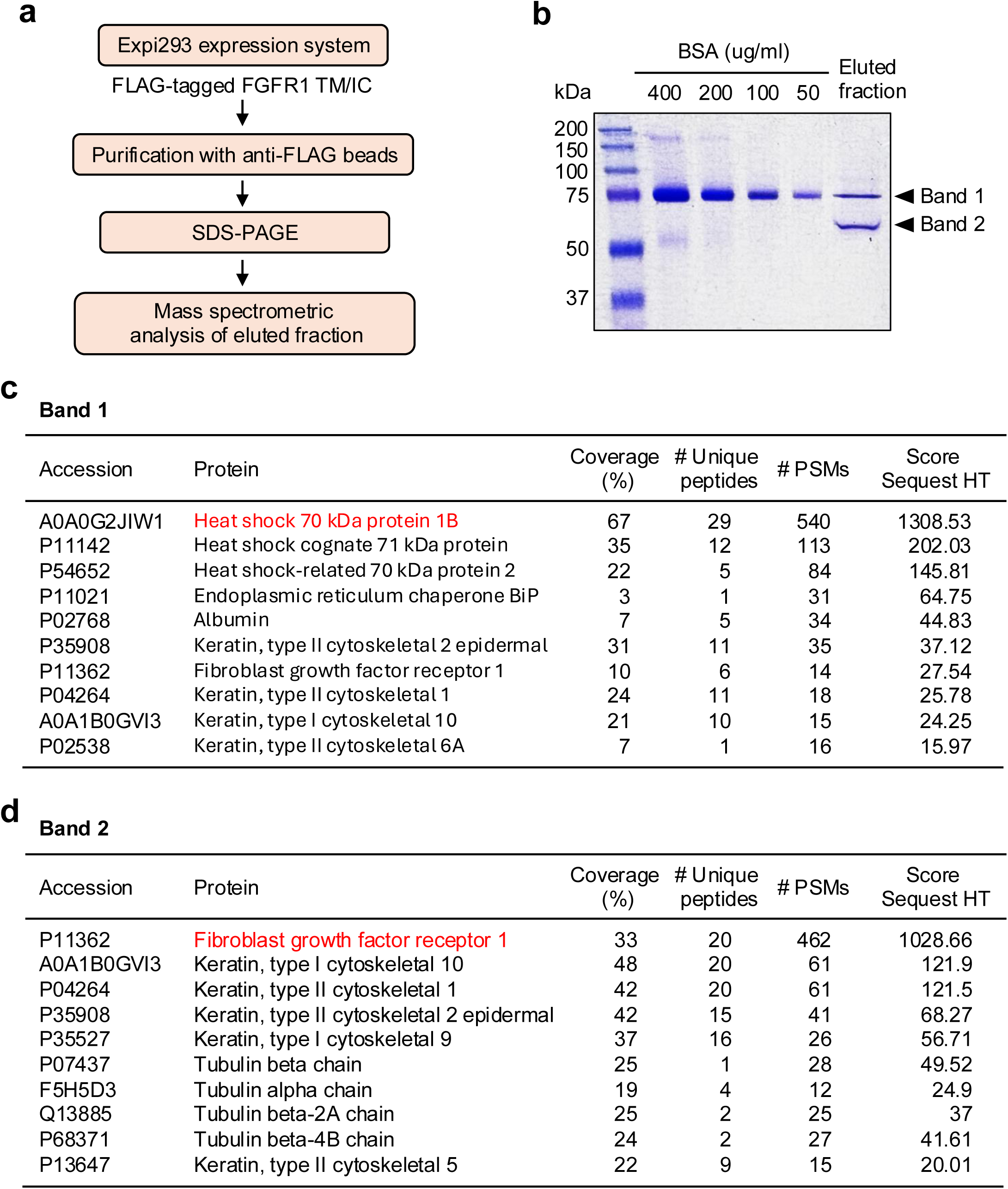
Purification and mass spectrometry analysis of FLAG-tagged FGFR1 TM/IC. **(a)** Experimental workflow of expression, purification and mass spectrometry analysis of recombinant FGFR1 TM/IC. **(b)** SDS–PAGE of eluted fraction which shows two major bands. **(c, d)** Proteins identified in Band 1 (**c**) and Band 2 (**d**). Mass spectrometry identified HSP70 family proteins in Band 1 and FGFR1 as the predominant protein in Band 2.

## Supplementary Movie legends

**Supplementary Movie 1. HS-AFM videos of three representative empty nanodiscs on an AP-mica surface.** Scan area, 80 × 64 nm^2^; image size, 100 × 80 pixels (acquired in HS-AFM observations); frame rate, 3.3 fps.

**Supplementary Movie 2. HS-AFM videos of three representative FGFR1 WT TM/IC reconstituted into nanodiscs on an AP-mica surface.** Magenta arrows indicate the KD. Scan area, 80 × 64 nm^2^; image size, 100 × 80 pixels (acquired in HS-AFM observations); frame rate, 3.3 fps.

**Supplementary Movie 3. HS-AFM videos of six representative FRS2α molecules on a bare mica surface.** White arrows indicate individual FRS2α molecules. Scan areas, 200 × 200, 120 × 96, and 80 × 64 nm^2^; image sizes, 180 × 180, 120 × 96, and 200 × 80 pixels (acquired in HS-AFM observations); frame rate, 1.0, and 3.3 fps.

**Supplementary Movie 4. HS-AFM videos of three representative FGFR1 WT TM/IC with FRS2α reconstituted into nanodiscs on an AP-mica surface.** Magenta and white arrows indicate the KD and FRS2α, respectively. Scan area, 80 × 64 nm^2^; image sizes, 100 × 80 and 200 × 80 pixels (acquired in HS-AFM observations); frame rate, 3.3 fps.

**Supplementary Movie 5. HS-AFM videos of three representative FGFR1 TM/IC K514A reconstituted into nanodiscs on an AP-mica surface.** Magenta arrows indicate the kinase domains. Scan area, 80 × 64 nm^2^; image size, 100 × 80 pixels (acquired in HS-AFM observations); frame rate, 3.3 fps.

**Supplementary Movie 6. HS-AFM videos of three representative FGFR1 TM/IC WT with 10 nM Futibatinib reconstituted into nanodisc on an AP-mica surface.** Magenta arrows indicate the kinase domains. Scan area, 80 × 64 nm^2^; image sizes, 100 × 80 pixels (acquired in HS-AFM observations); frame rate, 3.3 fps.

## Notes

### Competing Interest Statement

The authors have declared no competing interest.

## References

1 Lemmon, M. A. & Schlessinger, J. Cell signaling by receptor tyrosine kinases. Cell 141, 1117–1134, doi:10.1016/j.cell.2010.06.011 (2010).

2 Blume-Jensen, P. & Hunter, T. Oncogenic kinase signalling. Nature 411, 355–365, doi:10.1038/35077225 (2001).

3 Mohammadi, M., Schlessinger, J. & Hubbard, S. R. Structure of the FGF receptor tyrosine kinase domain reveals a novel autoinhibitory mechanism. Cell 86, 577–587, doi:10.1016/s0092-8674(00)80131-2 (1996).

4 Mohammadi, M. et al. Structures of the tyrosine kinase domain of fibroblast growth factor receptor in complex with inhibitors. Science 276, 955–960, doi:10.1126/science.276.5314.955 (1997).

5 Zhang, J., Yang, P. L. & Gray, N. S. Targeting cancer with small molecule kinase inhibitors. Nat Rev Cancer 9, 28–39, doi:10.1038/nrc2559 (2009).

6 Rudolph, J. H., KP.; Dar, AC. Contemporary design of small-molecule kinase modulators: orthosteric, allosteric and induced-proximity strategies. Nat Rev Cancer 25, 595–618 (2026).

7 Wright, P. E. & Dyson, H. J. Intrinsically disordered proteins in cellular signalling and regulation. Nat Rev Mol Cell Biol 16, 18–29, doi:10.1038/nrm3920 (2015).

8 Holehouse, A. S. & Kragelund, B. B. The molecular basis for cellular function of intrinsically disordered protein regions. Nat Rev Mol Cell Biol 25, 187–211, doi:10.1038/s41580-023-00673-0 (2024).

9 Okuda, M. & Nishimura, Y. Real-time and simultaneous monitoring of the phosphorylation and enhanced interaction of p53 and XPC acidic domains with the TFIIH p62 subunit. Oncogenesis 4, e150, doi:10.1038/oncsis.2015.13 (2015).

10 Pinet, L., Assrir, N. & van Heijenoort, C. Expanding the Disorder-Function Paradigm in the C-Terminal Tails of Erbbs. Biomolecules 11, doi:10.3390/biom11111690 (2021).

11 Kot, E. F. et al. Intrinsically disordered regions couple the ligand binding and kinase activation of Trk neurotrophin receptors. iScience 25, 104348, doi:10.1016/j.isci.2022.104348 (2022).

12 Schlessinger, J. Common and distinct elements in cellular signaling via EGF and FGF receptors. Science 306, 1506–1507, doi:10.1126/science.1105396 (2004).

13 Yarden, Y. & Sliwkowski, M. X. Untangling the ErbB signalling network. Nat Rev Mol Cell Biol 2, 127–137, doi:10.1038/35052073 (2001).

14 Pawson, T. & Scott, J. D. Signaling through scaffold, anchoring, and adaptor proteins. Science 278, 2075–2080, doi:10.1126/science.278.5346.2075 (1997).

15 Turner, N. & Grose, R. Fibroblast growth factor signalling: from development to cancer. Nat Rev Cancer 10, 116–129, doi:10.1038/nrc2780 (2010).

16 Babina, I. S. & Turner, N. C. Advances and challenges in targeting FGFR signalling in cancer. Nat Rev Cancer 17, 318–332, doi:10.1038/nrc.2017.8 (2017).

17 Gotoh, N. Regulation of growth factor signaling by FRS2 family docking/scaffold adaptor proteins. Cancer Sci 99, 1319–1325, doi:10.1111/j.1349-7006.2008.00840.x (2008).

18 Eswarakumar, V. P., Lax, I. & Schlessinger, J. Cellular signaling by fibroblast growth factor receptors. Cytokine Growth Factor Rev 16, 139–149, doi:10.1016/j.cytogfr.2005.01.001 (2005).

19 Kouhara, H. et al. A lipid-anchored Grb2-binding protein that links FGF-receptor activation to the Ras/MAPK signaling pathway. Cell 89, 693–702, doi:10.1016/s0092-8674(00)80252-4 (1997).

20 Ando, T., Uchihashi, T. & Scheuring, S. Filming biomolecular processes by high-speed atomic force microscopy. Chem Rev 114, 3120–3188, doi:10.1021/cr4003837 (2014).

21 Sumino, A. et al. High-Speed Atomic Force Microscopy Reveals Fluctuations and Dimer Splitting of the N-Terminal Domain of GluA2 Ionotropic Glutamate Receptor-Auxiliary Subunit Complex. ACS Nano 18, 25018–25035, doi:10.1021/acsnano.4c06295 (2024).

22 Kodera, N. et al. Structural and dynamics analysis of intrinsically disordered proteins by high-speed atomic force microscopy. Nat Nanotechnol 16, 181–189, doi:10.1038/s41565-020-00798-9 (2021).

23 Goyal, L. et al. Futibatinib for FGFR2-Rearranged Intrahepatic Cholangiocarcinoma. N Engl J Med 388, 228–239, doi:10.1056/NEJMoa2206834 (2023).

24 Abramson, J. et al. Accurate structure prediction of biomolecular interactions with AlphaFold 3. Nature 630, 493–500, doi:10.1038/s41586-024-07487-w (2024).

25 Mohammadi, M. et al. Identification of six novel autophosphorylation sites on fibroblast growth factor receptor 1 and elucidation of their importance in receptor activation and signal transduction. Mol Cell Biol 16, 977–989, doi:10.1128/MCB.16.3.977 (1996).

26 Goyal, L. et al. TAS-120 Overcomes Resistance to ATP-Competitive FGFR Inhibitors in Patients with FGFR2 Fusion-Positive Intrahepatic Cholangiocarcinoma. Cancer Discov 9, 1064–1079, doi:10.1158/2159-8290.CD-19-0182 (2019).

27 Bartha, A. & Gyorffy, B. TNMplot: An enhanced platform for pharmacological target identification through cross-stage and pan-cancer gene expression analysis. Br J Pharmacol 183, 2648–2659, doi:10.1111/bph.70390 (2026).

28 Cerami, E. et al. The cBio cancer genomics portal: an open platform for exploring multidimensional cancer genomics data. Cancer Discov 2, 401–404, doi:10.1158/2159-8290.CD-12-0095 (2012).

29 Gao, J. et al. Integrative analysis of complex cancer genomics and clinical profiles using the cBioPortal. Sci Signal 6, pl1, doi:10.1126/scisignal.2004088 (2013).

30 de Bruijn, I. et al. Analysis and Visualization of Longitudinal Genomic and Clinical Data from the AACR Project GENIE Biopharma Collaborative in cBioPortal. Cancer Res 83, 3861–3867, doi:10.1158/0008-5472.CAN-23-0816 (2023).

31 Wybenga-Groot, L. E. et al. Structural basis for autoinhibition of the Ephb2 receptor tyrosine kinase by the unphosphorylated juxtamembrane region. Cell 106, 745–757, doi:10.1016/s0092-8674(01)00496-2 (2001).

32 Hubbard, S. R. Juxtamembrane autoinhibition in receptor tyrosine kinases. Nat Rev Mol Cell Biol 5, 464–471, doi:10.1038/nrm1399 (2004).

33 Landau, M., Fleishman, S. J. & Ben-Tal, N. A putative mechanism for downregulation of the catalytic activity of the EGF receptor via direct contact between its kinase and C- terminal domains. Structure 12, 2265–2275, doi:10.1016/j.str.2004.10.006 (2004).

34 Lin, C. C. et al. The combined action of the intracellular regions regulates FGFR2 kinase activity. Commun Biol 6, 728, doi:10.1038/s42003-023-05112-6 (2023).

35 Lin, C. W. et al. A two-component protein condensate of the EGFR cytoplasmic tail and Grb2 regulates Ras activation by SOS at the membrane. Proc Natl Acad Sci U S A 119, e2122531119, doi:10.1073/pnas.2122531119 (2022).

## References

36 Schindelin, J. et al. Fiji: an open-source platform for biological-image analysis. Nat Methods 9, 676–682, doi:10.1038/nmeth.2019 (2012).

37 Tseng, Q. et al. A new micropatterning method of soft substrates reveals that different tumorigenic signals can promote or reduce cell contraction levels. Lab Chip 11, 2231–2240, doi:10.1039/c0lc00641f (2011).

38 Meijering, E., Dzyubachyk, O. & Smal, I. Methods for cell and particle tracking. Methods Enzymol 504, 183–200, doi:10.1016/B978-0-12-391857-4.00009-4 (2012).

39 Ishida, T. & Kinoshita, K. PrDOS: prediction of disordered protein regions from amino acid sequence. Nucleic Acids Res 35, W460–464, doi:10.1093/nar/gkm363 (2007).

40 Pettersen, E. F. et al. UCSF ChimeraX: Structure visualization for researchers, educators, and developers. Protein Sci 30, 70–82, doi:10.1002/pro.3943 (2021).

41 al., A. M. e. GROMACS: High performance molecular simulations through multi-level parallelism from laptops to supercomputers. SoftwareX (2015).

42 Wu, E. L. et al. CHARMM-GUI Membrane Builder toward realistic biological membrane simulations. J Comput Chem 35, 1997–2004, doi:10.1002/jcc.23702 (2014).

43 Lomize, M. A., Pogozheva, I. D., Joo, H., Mosberg, H. I. & Lomize, A. L. OPM database and PPM web server: resources for positioning of proteins in membranes. Nucleic Acids Res 40, D370–376, doi:10.1093/nar/gkr703 (2012).

44 Tian, C. et al. ff19SB: Amino-Acid-Specific Protein Backbone Parameters Trained against Quantum Mechanics Energy Surfaces in Solution. J Chem Theory Comput 16, 528–552, doi:10.1021/acs.jctc.9b00591 (2020).

45 Dickson, C. J., Walker, R. C. & Gould, I. R. Lipid21: Complex Lipid Membrane Simulations with AMBER. J Chem Theory Comput 18, 1726–1736, doi:10.1021/acs.jctc.1c01217 (2022).

46 al, J. W. e. Comparison of simple potential functions for simulating liquid water.. J Chem Phys (1983).

47 Joung, I. S. & Cheatham, T. E., 3rd. Determination of alkali and halide monovalent ion parameters for use in explicitly solvated biomolecular simulations. J Phys Chem B 112, 9020–9041, doi:10.1021/jp8001614 (2008).

48 Lee, J. et al. CHARMM-GUI supports the Amber force fields. J Chem Phys 153, 035103, doi:10.1063/5.0012280 (2020).

49 al, E. U. e. A smooth particle mesh Ewald method. J Chem Phys 103, 8577 (1995).

50 Bussi, G., Donadio, D. & Parrinello, M. Canonical sampling through velocity rescaling. J Chem Phys 126, 014101, doi:10.1063/1.2408420 (2007).

51 Bernetti, M. & Bussi, G. Pressure control using stochastic cell rescaling. J Chem Phys 153, 114107, doi:10.1063/5.0020514 (2020).

52 Hess B, e. a. LINCS: A linear constraint solver for molecular simulations.. J Comput Chem 18, 1463–1472 (1997).

53 Miyamoto S, K. P. SETTLE: An analytical version of the SHAKE and RATTLE algorithm for rigid water models.. 13, 952–962 (1992).

